# Gut microbiota and holobiont metabolome composition of the Medaka fish (*Oryzias latipes*) are affected by a short exposure to the cyanobacterium *Microcystis aeruginosa*

**DOI:** 10.1101/2022.07.08.499308

**Authors:** Pierre Foucault, Alison Gallet, Charlotte Duval, Benjamin Marie, Sébastien Duperron

**Affiliations:** UMR7245 Molécules de Communication et Adaptation des Micro-organismes, Muséum National d’Histoire Naturelle, CNRS, Paris, France

**Keywords:** Microbiota, Ecotoxicology, Multi-omics, Time-series, Cyanobacteria, Holobiont

## Abstract

Blooms of toxic cyanobacteria are a common stress encountered by aquatic fauna. Evidence indicates that long-lasting blooms affect fauna-associated microbiota. Because of their multiple roles, host-associated microbes are nowadays considered relevant to ecotoxicology, yet the respective timing of microbiota versus functional changes in holobionts response needs to be clarified. The response of gut microbiota and holobiont’s metabolome to exposure to a dense culture of *Microcystis aeruginosa* was investigated as a microcosm-simulated bloom in the model fish species *Oryzias latipes* (medaka). Both gut microbiota and gut metabolome displayed significant composition changes after only 2 days of exposure. A dominant symbiont, member of the Firmicutes, plummeted whereas various genera of Proteobacteria and Actinobacteriota increased in relative abundance. Changes in microbiota composition occurred earlier and faster compared to metabolome composition, suggesting that the microbiota drives the holobiont’s response. Liver and muscle metabolome were much less affected than guts, supporting that gut and associated microbiota are in the front row upon exposure. This study highlights that even short cyanobacterial blooms, that are increasingly frequent, trigger changes in microbiota composition and holobiont metabolome. It emphasizes the relevance of multi-omics approaches to explore organism’s response to an ecotoxicological stress.

**Graphical abstract:** 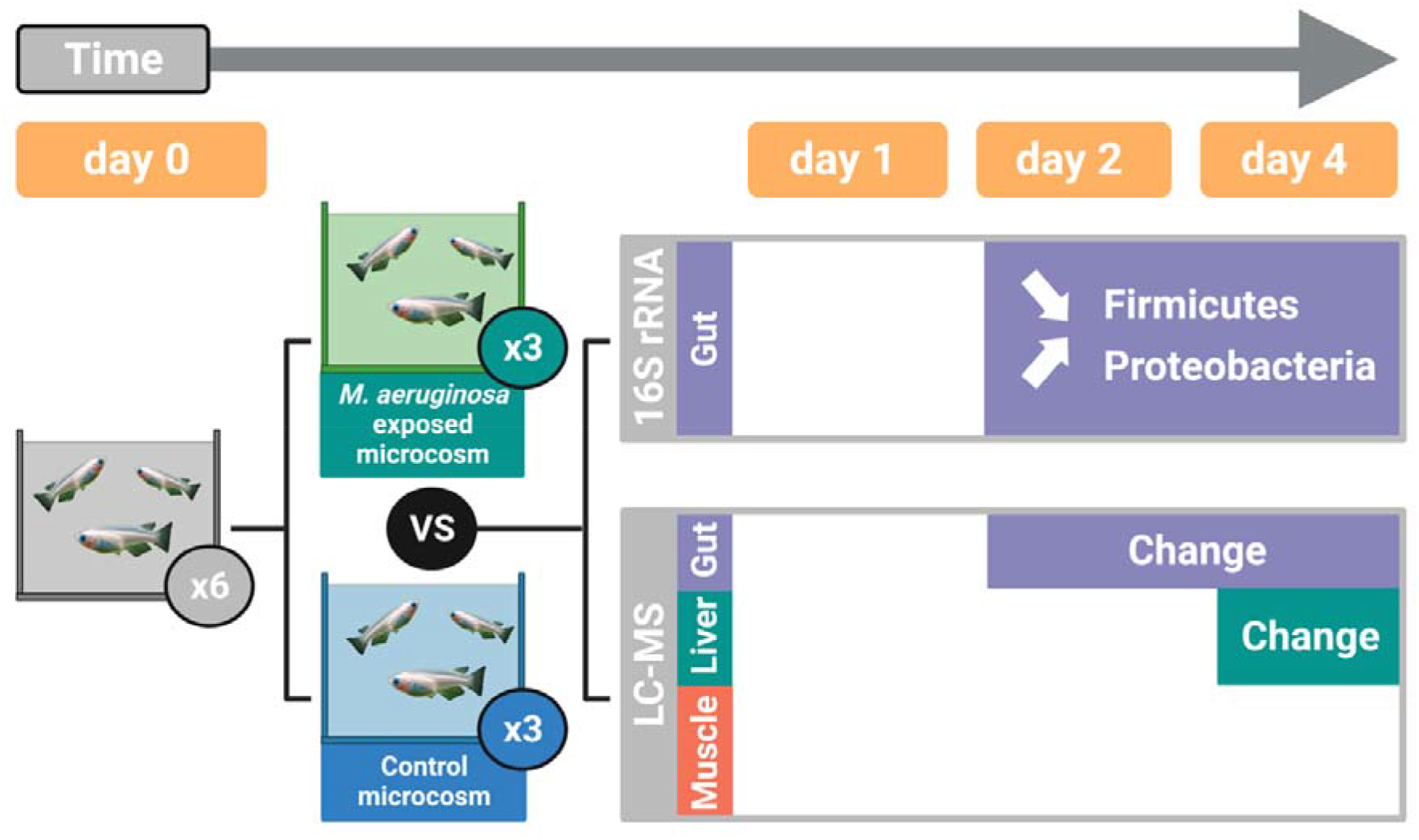

**Highlights:**

- A 2-day exposure to a simulated *M. aeruginosa* bloom is enough to sharply decrease the Firmicute/Proteobacteria ratio in the gut of *Oryzias latipes* fish.
- The exposure induced changes in metabolome composition after 2 days in the gut and 4 days in the liver.
- The gut bacterial microbiota response occurred faster than metabolome’s; we hypothesize that changes in gut microbiota may drive the gut metabolome compositional changes.

## 1. Introduction

One of the most common yet intense stress encountered by fauna in ponds and lakes is the occurrence of algal bloom involving noxious cyanobacteria, which are increasing in relation to anthropogenic effects^1^. Adverse effects of blooms, ranging from specific organ toxicity (e.g. hepatotoxicity and cardiotoxicity) to reproduction alteration, are increasingly documented in various animal models^2–5^. On the other hand, effects of blooms on associated microbiota are barely known. Beyond their role in holobiont development and functioning (nutrition, immunity, protection, behavior…), host-associated microbial communities are now considered fully relevant to ecotoxicology^6–9^. Indeed, communities are often located at the interface between a host and its environment. They react to and interact with contaminants, with outcomes ranging from inactivation to potentialization^10^. Because of their relative limited diversity of gut microorganisms, notably compared to endotherms, and of their patrimonial and economic value, teleost fish are particularly relevant vertebrate models for aquatic microbiome-aware ecotoxicology^7^. Among these, the medaka *Oryzias latipes* and the zebrafish *Danio rerio* are commonly employed for ecotoxicological studies.

Recently, 28-day exposure experiments conducted in microcosms on the medaka revealed that extracts of *Microcystis aeruginosa* containing microcystin-LR (MC-LR) among other bioactive metabolites could trigger dysbiosis in gut microbiota^11^. Since then, similar effects were shown in the zebrafish^12,13^. A recent study used microcosm-simulated *M. aeruginosa* blooms by exposing fish to high density of a cyanobacterial culture. Results revealed dose-dependent alterations of the composition of medaka gut bacterial communities and the holobiont metabolome, including a sharp decrease in abundance of a significant Firmicutes symbiont^14^. Altogether, these studies demonstrate that long exposures to cyanobacteria, together with their subsequent metabolites affect fish microbiota, and at least one of those studies also suggests an alteration of holobiont functions correlated with changes in microbiota composition^14^. In nature though, cyanobacterial blooms are often shorter than the typical 20-28 days exposure used in ecotoxicological studies and documenting the early dynamics of changes is the key to understanding holobiont response in natural relevant settings.

In this study, we investigated the early response of the medaka gut microbiota and organ metabolomes (gut, liver and muscle) to a 4-day exposure to high yet environmentally relevant density of *M. aeruginosa*, the main species responsible for blooms in temperate lakes and ponds. Compositions of the gut bacterial communities and gut, liver and muscle metabolomes are monitored using 16S rRNA gene sequencing and LC-MS metabolomics, respectively. Alteration dynamics of these compartments are then compared to highlight the key role of gut microbiota in the holobiont functional response. This study is the first attempt to address the initial short-term response of fish gut microbiota and holobiont metabolome to a simulated cyanobacterial bloom.

## 2. Material and methods

### 2.1 Exposure experiments

Experiments were performed in six aquaria (10-liters each, 3 control and 3 exposed, assigned randomly), each containing 15 specimens of 6-months old adult male medaka. Aquaria were stabilized for one month and fishes acclimatized for one week prior to exposure. At day 0, nine fish in total were sampled randomly across the six aquaria and 25 mL of water from each aquarium were pooled as controls. Fish were then exposed for 4 days to water, or water containing a strong but environmentally relevant density of the live non-axenic monoclonal *M. aeruginosa* strain PMC 728.11^15^ of the Paris Museum Collection^16^ to simulate a bloom (100 *µ*g.L^-1^ Chl *a*). The strain was cultured accordingly to anterior publication^11,14^ (details in the supplementary information). *M. aeruginosa* concentrations were estimated after performing Chlorophyll *a* extractions and absorbance measurements as a proxy using a spectrophotometer^17^ (Cary 60 UV-Vis, Agilent). Culture was sampled for DNA (1 mL) and metabolome (50 mL) analyses on days 0 and 2. The *M. aeruginosa* level in exposed aquaria was adjusted on days 0 and 2 to maintain exposure level. Three fish per aquarium were sampled on days 1 and 2 and four per aquarium at day 4 (3 in one aquarium), as well as water samples.

Water parameters were monitored on days 0, 2 and 4 (pH, temperature, conductivity, nitrates and nitrites), feces were removed daily by aspiration, and half of the water was replaced with freshwater (2/3 osmosis (RiOs 5, Merck Millipore) and 1/3 filtered) containing or not, adjusted amount of *Microcystis* cells. Fish were exposed to constant temperature (25.4 ±1 °C), pH (7.61 ±0.1) and conductivity (246 ±12.4 *µ*S.cm^-1^), to low levels of nitrates and nitrites (Table S1) to a 12h:12h light/dark cycle. They were fed twice daily (∼3-5% of the fish biomass per day) with Nutra HP 0.3 (Crude protein 57, Crude fat 17, N.F.E 7.5, Ash 10, Crude fiber 0.5, Phosphorus 1.7, Vitamins A, D3, E; Skretting, Norway). Total microcystines (MCs) levels were quantified in duplicates on days 0 and 4 (details in the supplementary information).

### 2.2 Metabolites extraction and characterization

Metabolites were extracted from flash-frozen dissected medaka guts, livers, muscles, and from lyophilized *M. aeruginosa* cultures. Mechanical extraction (GLH850 OMNI; 25 000 r.min^-1^; 30s) followed by sonication (Sonics Vibra-Cell VCX 13; 60% amplitude; 30s) were performed on weighted samples suspended in the extraction solvent (75-25% UHPLC methanol-water, 1 mL per 100 mg of tissue or per 10 mg of lyophilized culture, on ice). After centrifugation (10 min; 4 °C; 15,300 g), gut pellets were dried and used for subsequent DNA extraction^14^.

Supernatants containing metabolite extracts were analyzed by Ultra high-performance liquid chromatography (UHPLC, Elute Bruker) using a Polar Avance II 2,5 pore C18 column (300 *µ*L.min^-1^, Thermo) coupled with a high-resolution mass spectrometer (ESI-Qq-TOF, Compact Bruker) at 2 Hz speed on positive simple MS mode. Feature peak lists were generated from MS spectra within a retention time window of 1-15 minutes and a filtering of 5000 counts using MetaboScape 4.0 software (Bruker). The peak lists consisted of the area-under-the-peaks of extracted analytes from the three tissues (medaka’s gut, liver and muscles) sampled on days 0, 1, 2, 4 and the *M. aeruginosa* lyophilized culture. A log transformation was applied to metabolomics datasets. Principal Component Analysis (PCA) were performed using the mixOmics^18^ (v6.14.1) R package.

### 2.3 DNA extraction, sequencing and analysis of the V4-V5 region of bacterial 16S rRNA

DNA was extracted from the pellets of the guts and *M. aeruginosa* culture, and from 0.22 *µ*m filters for aquarium water. Extractions were performed using the ZymoBIOMICS DNA Mini-prep kit with a FastPrep 5G beat beater disruption (DNA Matrix; 4×30s; 6m.s^-1^) following manufacturer’s instructions. An extraction-blank control sample was also performed. The V4-V5 region of the 16S rRNA encoding gene was amplified using primers 515R and 926F^19^ and sequenced on an Illumina MiSeq 250×2 bp platform (Biomnigene, Besançon, France). Reads were deposited into the Sequence Read Archive (SRA) database (accession number PRJNA836730 (samples SRR19170691-SRR19170767; Table S2). Sequence analysis including primer removal and quality control was performed using the QIIME2-2021.2 pipeline^20^. Forward and reverse reads were trimmed at 250 and 200 bp, respectively. Amplicon Sequence Variants (ASVs) were obtained with DADA2 (default parameters) and affiliated with the SILVA 138-99 database. Diversity metrics were computed with the phyloseq^21^ (v1.34.0) R package. Statistical analyses were performed using R packages vegan^22^ (v2.5-7) and RVAideMemoire^23^ (v0.9-79). All values are displayed as median ±standard deviation.

### 2.4 MEBA analysis

A Multivariate Empirical BayesianAnalysis^24,25^ (MEBA) was performed to discriminate differentially abundant taxa through time and between treatments. Relative abundance tables at the Phylum and Genus taxonomic levels were obtained from phyloseq and analyzed using the MEBA plugin from Metaboanalyst 5.0 (https://www.metaboanalyst.ca), with no data transformation and all by-default parameters. Taxa with a MEBA T^2^ score superior to 5 were consider discriminant and further statistically analyzed.

### 2.5 ASV-metabolites correlative networks

A correlative network analysis was performed with DIABLO^26^, a multi-omics framework created by the MixOmics team^18^, to analyze putative correlated dynamics between gut microbiota ASVs and gut metabolites. Briefly, the *block.plsda()* function performs a Pattern Latent Structure Discriminant Analysis which provides a reduced-dimension space with covariance-maximizing axes for each dataset (named “block”). A correlation score for the given blocks was computed with the *plotDiablo()* function.

### 2.6 Comparative dynamics of microbiota and metabolome

The composition change dynamics in microbiota versus metabolome were investigated by comparing trajectories of centroids using a newly developed method: MOTA (Multivariate Omics Trajectory Analysis). Microbiota and metabolome datasets were log transformed and analyzed separately but in a similar way. Distances between PCAs’ centroid coordinates were computed to create a trajectory between each sampling day for the two treatments and displayed as the fraction of the total length achieved at each day (from 0% to 100% between day 0 to 4). Trajectories were plotted for the 16S rRNA versus metabolomic data, allowing comparison of their respective dynamics in the two treatments (details in the supplementary information).

## 3. Results and discussion

Concentrations of *M. aeruginosa* varied between 31 and 126 *µ*g.L^-1^ Chl *a* in exposed aquaria, well above values recently documented to alter microbiota composition after a 28-day exposure^14^, and are thus appropriate to investigate short-term effect and early response of the holobiont. In these aquaria, MCs levels were high at day 1 and increased at day 4 (12 ±5 to 21 ±9 *µ*g eq. L^-1^ MC-LR, respectively; Table S1). These conditions, above values reported from extracts used previously^11^, are expected to produce major changes in a 14-day exposure and are thus appropriate to evaluate short term effects.

### 3.1 Rapid shift in community compositions during exposure to Microcystis aeruginosa

Analysis of bacterial community compositions based on the V4-V5 region of 16S rRNA-encoding gene from 67 fish guts, 7 waters, 2 *M. aeruginosa* cultures and 1 extraction blank samples yielded 3,702,081 raw reads, of which 73.4% were retained after sequence assembly, denoising and chimera removal. Samples displayed 21,134 to 59,009 reads (31,638 ±7,000), except the extraction blank (520 reads; Table S3). After taxonomic assignment and removal of eukaryotes, mitochondria and chloroplasts, sequences clustered into 936 ASVs. Of these, 856 were considered as abundant (they represented at least 1% of reads in at least 1 sample). Rarefaction curves reached saturation confirming that our sequencing effort was sufficient to describe most of the bacterial diversity (not shown). Alpha diversity indices did not indicate major changes among microcosm compartments (culture, water and fish guts) or between dates and treatments (Fig S1; Table S4). Although bacterial ASVs richness was higher in fish guts than other compartments and bacterial ASVs richness and evenness increased during the experiment in both exposed and unexposed fish guts, differences were not statistically significant.

Community compositions were significantly different in fish guts compared to water and culture samples (Unweighted UniFrac; Permanova *p*<0.05; Table S5). Among fish guts, composition differences were significant (Unweighted UniFrac; Permanova *p*<0.05; Table S5). Visually, samples from different time points are scattered along the first PCoA axis and the two treatments are well separated on the second axis of the PCoA (Fig 1B). Indeed, the microbiota of fish exposed to high density of *M. aeruginosa*, or non-exposed, displayed compositions different from d0 at both d2 and d4. In addition, significant differences between the two treatments were observed at d2 and d4 (Unweighted UniFrac; Pairwise.Permanova *p*<0.05; Table S5), indicating that gut community compositions were affected by exposure duration (day 0 versus 2 and 4) and treatment (water versus *M. aeruginosa* exposed). Intra-group variances were not significantly different, allowing group comparison (Permdisp *p*>0.05; Table S5).

**Fig. 1:**
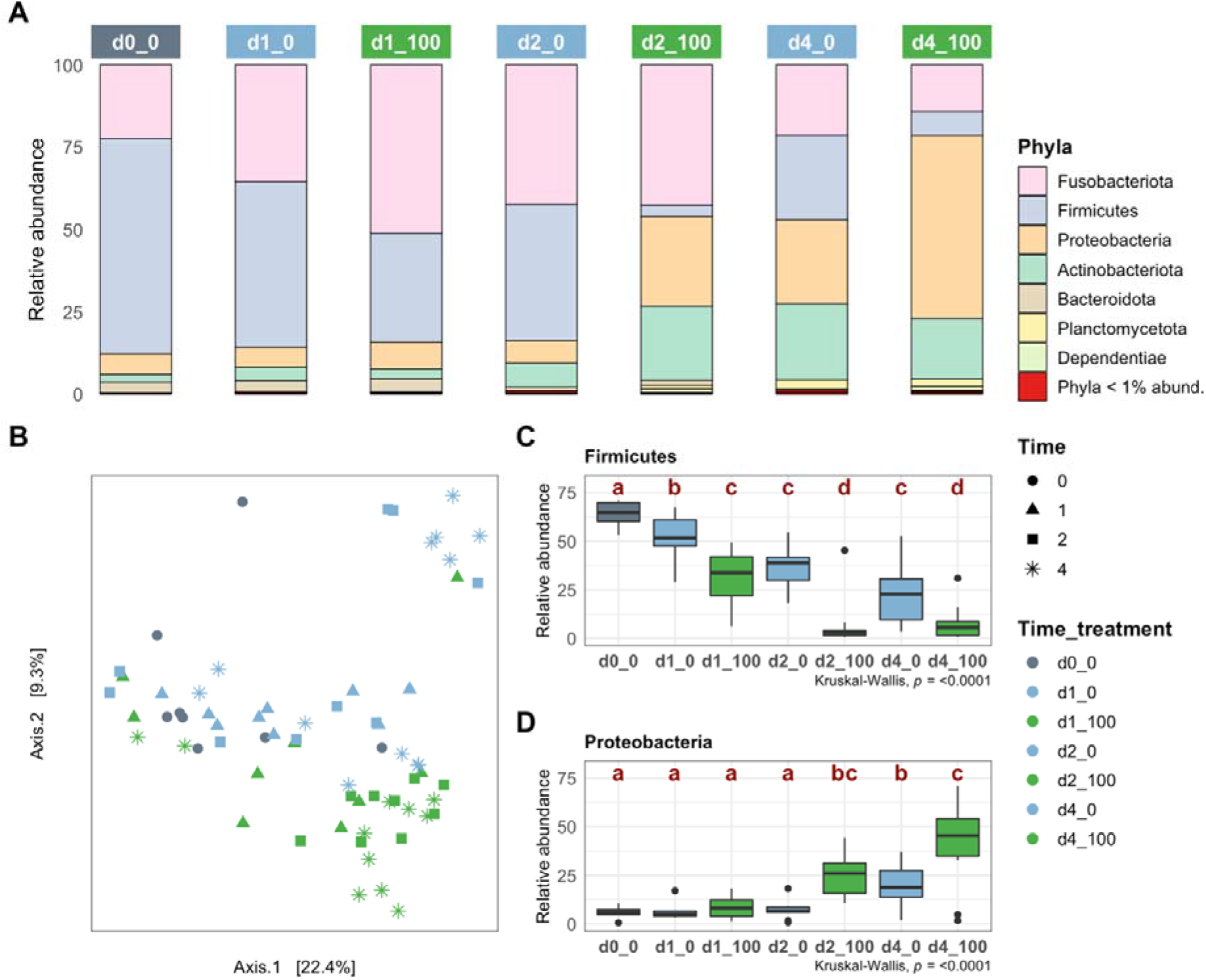
Composition and diversity of fish gut bacterial microbiota among time and treatment. **A:** Gut microbiota composition at the phylum level (median values of samples belonging the same time_treatment group). **B:** Principal Coordinates Analysis (Unweighted UniFrac distance). **C-D:** Relative abundance variations over time in both treatments, observed in Firmicutes (**C**) and Proteobacteria (**D**); letters refer to Benjamini-Hochberg (BH) adjusted Wilcoxon post-hoc test significance.

Firmicutes was the dominant phylum in the fish gut bacterial communities at day 0 (64.7 %±6 of reads; Fig. 1A) and was discriminant among time and treatment (MEBA T^2^ score>5; Table S6). In *M. aeruginosa*-exposed fish, Firmicutes decreased significantly between day 0 and 1 (64.7 ±6% to 33.7 ±14.4%; Wilcoxon *p*<0.05; Fig 1C; Table S6), then day 2 (to 2.91 ±14.24 %; Wilcoxon *p*<0.05; Table S6) after when then remained stable at day 4 (Wilcoxon *p*>0.05; Table S6). Firmicutes decreased less dramatically in unexposed fish. Significant decreases were observed on days 2 and 4 (38.8 ±12.3% and 22.78 ±16.49% respectively; Wilcoxon *p*<0.05; Table S6). The gap between exposed and unexposed fish on days 2 and 4 was significant, supporting that the treatment itself further decreased Firmicutes abundance on top of the effect observed in unexposed fish. Among discriminant taxa, this decreasing pattern was only observed with Firmicutes and its main genus *ZOR0006* (Table S6). A similar decrease of *ZOR0006* was observed after 28 days exposure to even moderate levels of *M. aeruginosa*^14^. Based on its genome content^14^, *ZOR0006*, a dominant resident in guts of healthy medaka^14^, can perform lactate pyruvate interconversion, a function essential to the gut homeostasis as lactate generally inhibits the growth of most pathogens while pyruvate stimulates it^27–29^. Lactate can also be involved in the repair of the gut epithelium, yet high levels can also be associated with inflammatory bowel disease in humans^30,31^. The decrease of Firmicutes has been considered an indicator of dysbiosis^24^, here suggesting that exposure to *M. aeruginosa* triggers a strong dysbiosis as early as 2 days. The moderate decrease perceived in unexposed specimens could be a consequence of the stress associated with specimen handling^32^.

Contrariwise, Proteobacteria were also discriminant (MEBA T^2^ score>5), but their abundances increased significantly from day 2 in *M. aeruginosa*-exposed fish. It was even more intense at day 4 (18.70 ±11.34 to 45.21 ±21.65%; Fig 1D; Wilcoxon *p*<0.05; Table S6), while it was significant only at day 4 for unexposed fish (Wilcoxon *p*<0.05; Table S6). As for the decreasing pattern, the gap between the unexposed and exposed samples on days 2 and 4 was significant (Wilcoxon *p*<0.05; Table S6), indicating that the treatment influenced the abundance of the Proteobacteria on top of the effect observed in unexposed fish. Four discriminant genera (*Xanthobacter, Reyranella* and *Devosia* belonging to Proteobacteria and *Nocardioides*, phylum Actinobacteriota) displayed a similar pattern (MEBA T^2^ score>5; Table S6). Other discriminant taxa abundances were not significantly altered between treatments on both days 2 and 4 (Table S6).

### 3.2 Exposure to Microcystis aeruginosa induces changes mostly in gut metabolome composition

Mass spectrometry distinguishes 921, 2,521 and 4,190 metabolites present in gut, liver, and muscle samples, respectively. The PCA analysis visually separates gut metabolomes of fish exposed to *M. aeruginosa* for 2 and 4 days, away from other samples (Fig. 2A). Metabolome compositions were significantly altered on days 2 and 4 for exposed fish and, to a lesser extent, unexposed fish (Permanova *p*<0.05; Table S5). The metabolite compositions between treatments on days 2 and 4 was different (Permanova *p*<0.05; Table S5), suggesting an additional effect of the treatment to the changes observed in unexposed fish. This assumption is further supported by the observation that 574 of the 916 observed in gut metabolites displayed differential abundances among dates in exposed specimens versus only 152 in unexposed fish (Anova *p*<0.05; Fig S2A, B). Liver metabolome composition is significantly different between day 4 and other dates, and at this day between treatments, suggesting an effect of both the duration of the experiment and the *M. aeruginosa* exposure after 4 days (Permanova *p*<0.05; Table S5). Intra-group variances were not significantly different, allowing group comparison (Permdisp *p*>0.05; Table S5). Finally, composition of muscle metabolomes did not display any significant variation (Permanova *p*>0.05; Table S5). Thus, the gut functional status appears altered earlier than the livers, while the muscles functional status is not affected. This is not surprising given that cyanobacteria enter fish through oral ingestion and their bio-active metabolites transfer through the intestine, which could buffer their effects before other organs are affected, and thus protect the host during short blooms^4,33^.

**Fig. 2:**
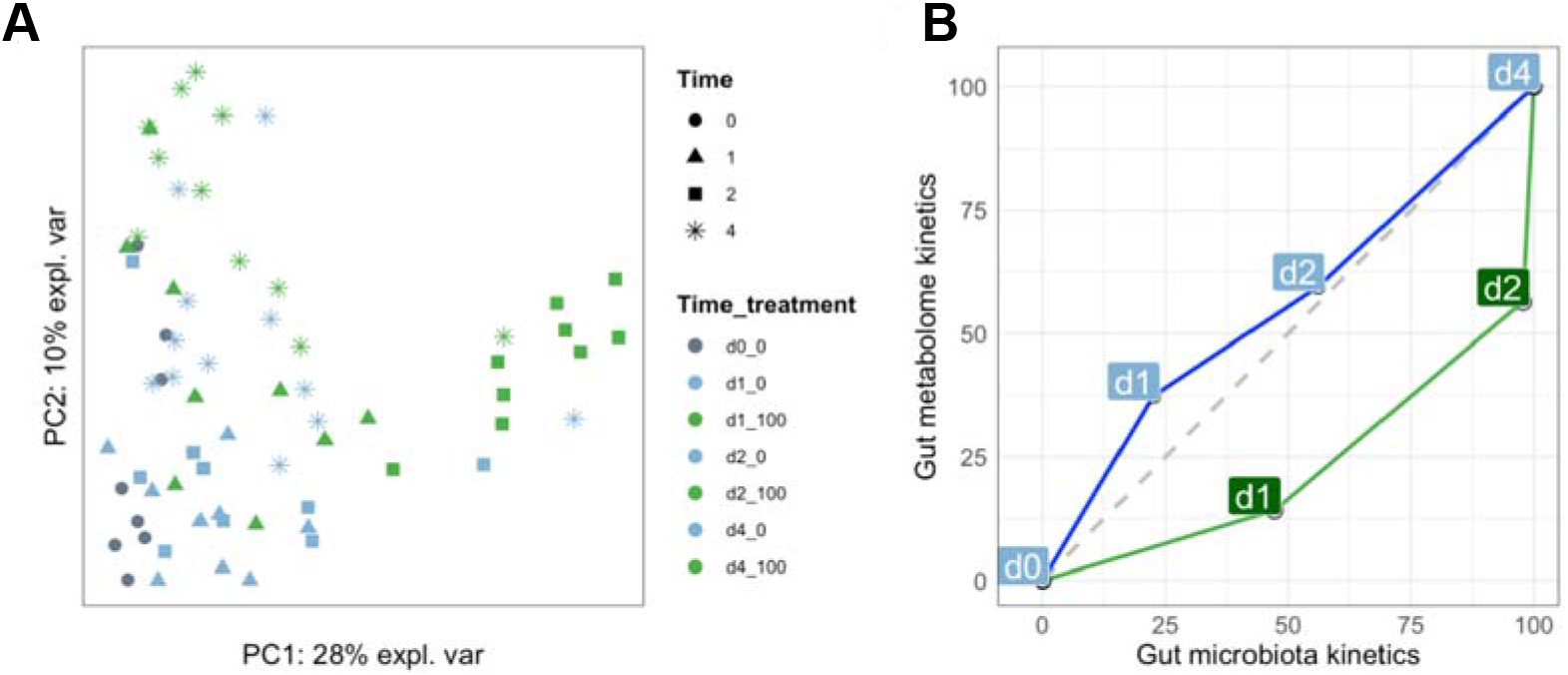
Dissimilarity between fish gut metabolome samples among time and treatment and composition change dynamic comparison between gut microbiota and metabolome. **A:** Principal Component Analysis of gut metabolite compositions of exposed and unexposed fish over time. **B:** Trajectory of the composition change in microbiota versus metabolome expressed as percentages of the total trajectory achieved on days 0, 1, 2 and 4 on each axis.

### 3.3 Do changes in gut bacterial communities drive functional changes in the holobiont?

In unexposed fish, the percentages of the total distance achieved by gut metabolome and microbiota compositions are similar on days 1 (22 and 37%) and 2 (56 and 59%), resulting in a trajectory that is close to the 1:1 diagonal (Fig. 2B), possibly reflecting an overall drift of the system due to the aforementioned handling stress^32^. The shape of the trajectory is different in exposed fish. Indeed, the gut metabolome achieved 14% of the total distance at day 1 versus 47% for the microbiota. At day 2, cumulative values are 56% for the metabolome and 98% for the microbiota, suggesting that most of the change ultimately observed in the microbiota composition is achieved after only 2 days. This holds true also when accounting for a higher percentage of the total explained variance (Fig S3). The trajectory thus shifts towards the gut microbiota axis for exposed samples, suggesting a faster response of the gut microbiota compared to the gut metabolome. The gut microbiota bacterial ASVs and gut metabolites datasets were well correlated at both day 2 and 4 (correlation score: 0.91). These results suggests that changes in microbiota might be preceding, and possibly driving observed metabolome changes, a hypothesis congruent with the localization of the gut microbes at the interface between host and its environment (digestive lumen), and thus their role as a primary barrier to contaminants^7,10^. Although it is tempting to infer causality here, further confirmation is needed since the gut metabolome contains both host and microbiota-derived metabolites, which could amplify the metabolome variations when bacterial communities change a lot.

## 4. Conclusion

Fish gut bacterial community composition and metabolome are affected within the first two days upon exposure to *M. aeruginosa*. Thus, even short cyanobacterial blooms can trigger drastic variations in bacterial phyla abundances, with consequences for the functioning of the gut. This way, iterative short bloom events could lead to major shifts in gut microbiota compositions and its associated functions, for example by progressively eliminating some sensitive resident symbiont lineages, exemplified here by the Firmicutes *ZOR0006*. Although sex-specific responses to perturbations have been observed in ecotoxicological^34^ and gut microbiota^35^ studies, only male fish were used here to first test the existence of a short-term response. Further experiments are now needed to address alterations in female fish. This study highlights the relevance of time-series exposure experiments and complementarity of omics approaches, using metabolomics as a proxy of the integrated functional response, to address short and long-term responses of holobionts to ecotoxicological stress, and the respective roles played by host and microbes.

## Author’s contributions

A.G., B.M. and S.D. conceived the study. A.G., C.D. and B.M. and S.D. conceived the experiment. P.F., A.G. and C.D. conducted the experiment. B.M. and S.D. took part in the experiment. P.F., A.G. and C.D. conducted molecular data processing. P.F. B.M., and S.D. analyzed data. P.F., B.M. and S.D. wrote the manuscript. All authors contributed and agreed on the contents.

## Notes

Experimental procedures were carried out in accordance with European legislation on animal experimentation (European Union Directive 2010/63/EU) and were approved for ethical contentment by an independent ethical council (CEEA Cuvier n°68) and authorized by the French government under reference number APAFiS#19316-2019032913284201 v1. Fish were anesthetized in 0.1% tricaine methanesulfonate (MS-222; Sigma, St. Louis, MO) buffered with 0.1% NaHCO3 prior to sacrifice.

## Acknowledgements

Kandiah Santhirakumar helped managing fish maintenance, Claude Yéprémian advised on cyanobacterial cultivation. We thank the Amagen platform for providing medaka fish, and the PtSMB platform of MNHN for metabolomics.

## Data availability

All R scripts are available on Github upon publication (https://github.com/PierreFoucault/). All microbial communities’ samples sequencing reads were deposited into the Sequence Read Archive (SRA) database (accession number PRJNA836730 (samples SRR19170691 to SRR19170767; Table S1).

## Supplementary information: supplementary Material and methods, Fig S1 and Fig S2

### Supplementary tables are summarized in a single supplementary file containing the following

Table S1: Parameters monitoring from the microcosm experiment Table S2: SRA accession numbers

Table S3: Sample IDs, raw reads counts, Sequencing depth and read filtering

Table S4: Gut microbiota samples median alpha-diversity indices and significativity group

Table S5: Gut microbiota samples alpha and beta diversity metrics; and gut, liver and muscles group comparison scores and significance levels

Table S6: Gut microbiota phylum and genera MEBA Hoteling T^2^, Kruskal Wallis and Wilcoxon post-hoc significativity levels (ns: not significant; *<0.05; **<0.01; ***<0.001)

Tables S7, S8 and Table S9: Gut, liver and muscle metabolites log-normalized count tables

