## Supplementary material and figures for "Gut microbiota and holobiont metabolome composition of the Medaka fish (*Oryzias latipes*) are affected by a short exposure to the cyanobacterium *Microcystis aeruginosa*"

**Supplementary information: supplementary Material and methods, Fig S1, S2 and S3**

**Supplementary Material and methods**

***M. aeruginosa PMC 728.11 strain culture***

To simulate a cyanobacterial bloom, we chose the non-axenic mono-clonal *M. aeruginosa* strain PMC 728.11 of the Paris Museum Collection, which was used in previous exposure experiments^1–3^ and which genome was sequenced^4^. This strain was isolated in 2011 near Valence, FRANCE and was since then kept in liquid culture in Z8 medium (composition presented in Hamlaoui et al.^5^) at 18 °C, and transferred to fresh culture medium every 4 months. Prior to the experimentation, 10 mL of the 728. 11 *M. aeruginosa* strain culture flask were deposit in a 250 mL Erlenmeyer and cultivated in 90 mL Z8 medium at 25 ±1 °C with a 16h:8h light/dark cycle (at 14 *µ*mol.m^-2^.s^-1^) for 2 weeks. Then, the 100 mL culture was transferred into a 500 mL Erlenmeyer with 300 mL of Z8 medium for 2 weeks. Finally, the 400 mL culture was transferred to a 2L bottle containing 1L of Z8 medium for 2 weeks.

***Microcystins quantification***

Microcystin-producing strains of *Microcystis* sp. can produce various microcystin congeners at the same time. The *M. aeruginosa* strain PMC 728.11, which genome have been already published^4^ mostly produced MC-LR variant, together with lesser amount of (eposyAdda5)MC-LR as previously described by Le Manach and colleagues^6^. In order to quantified the total amount of MC variant produced by this strain, we performed an specific ELISA test based on antibody targeting ADDA (the common part of all MC variants), that widely recognize all MC congeners, and the immunoreaction was expressed with regard to the intensity obtained with standard MC-LR, and then expressed in equivalent MC-LR (Microcystins-ADDA ELISA EPA 546, Eurofins Abraxis). Water samples were sonicated (Sonics Vibra-Cell VCX 130, 60% amplitude, 30s), thus the total MC content was determined on both intracellular and extracellular fraction in duplicates on days 0 and 4.

***MOTA analysis***

         Here, we present the new method MOTA (for Multivariate Omics Trajectory Analysis) to compare the dynamics of two omics datasets between two treatments (control and *M. aeruginosa* exposed) along a time-series (d0, d1, d2 and d4).

The method is based on multidimensional analysis (PCA). “Time_treatment” centroids are generated by averaging coordinates of each of the replicate samples within a “time_treatment” group. Within one treatment and one omic, distances between centroids (Euclidian norm) from consecutive time points are summed. The coordinates of these centroids are computed based on a variable number of axes in order to account for a given user set threshold of the cumulative percentage of explained variance. The cumulative percentage of total length achieved at each day (from 0% at d0 to 100% at d4) is displayed on the corresponding omics axis. Two omics can then be plotted on separate axes to compare their respective trajectories, in this case the 16S rRNA data for the gut microbiota versus the LC-MS data for the gut metabolome. Trajectories in the two treatments are plotted in blue (control) and green (*M. aeruginosa* exposed, Fig. S2).

The R script is available on GitHub:

<https://github.com/PierreFoucault/Medaka_MOTA/blob/859852840ad70e36d053681feb01d9556ef96d57/MOTA_PCA_gut_code.Rmd>

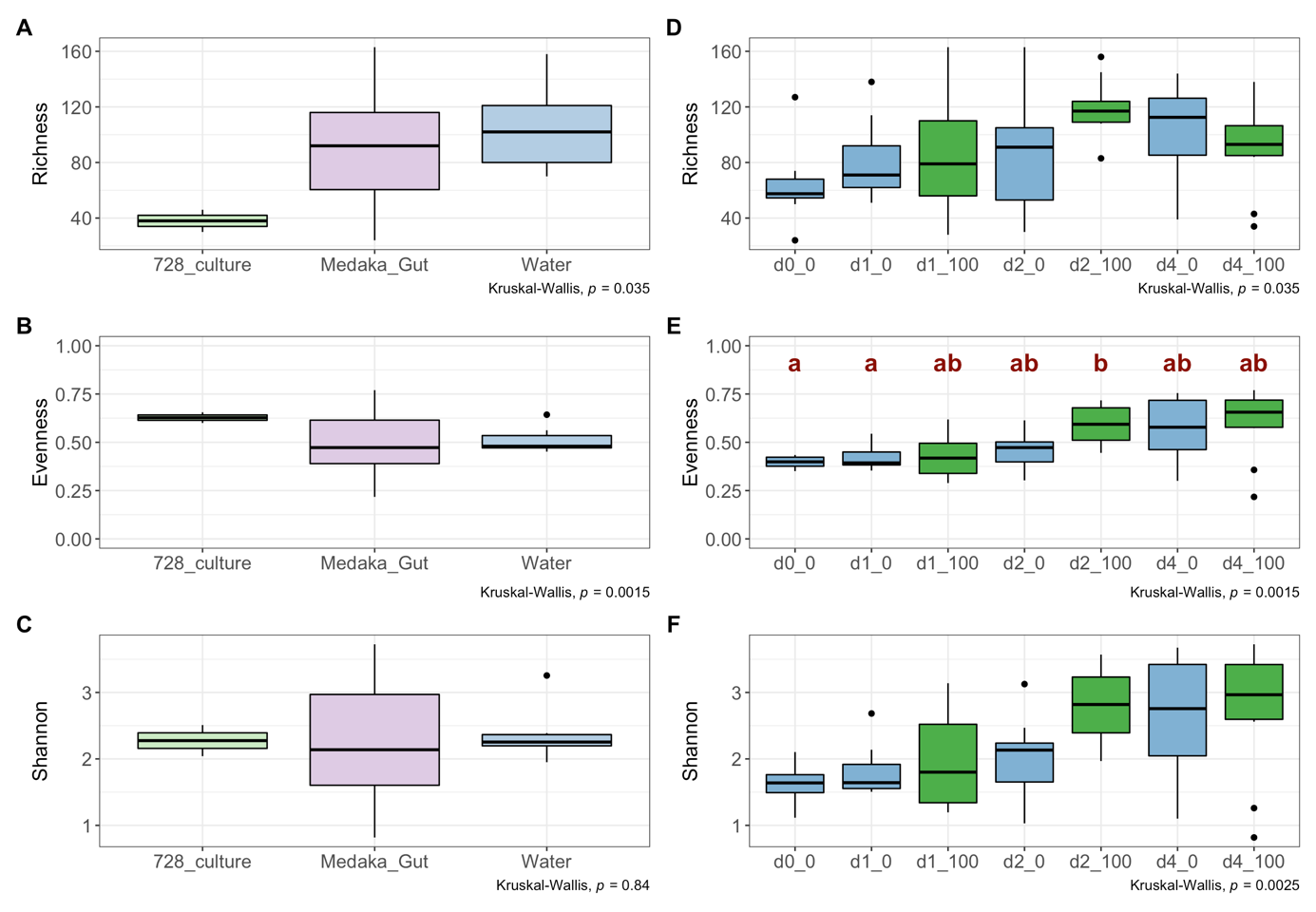

**Fig S1: Supplementary gut microbiota analysis figures.**

**A, B and C:** Bacterial ASVs richness (**A**), Pielou’s evenness (**B**) and shannon (**C**) indices of the 3 microcosm compartments (*M. aeruginosa* culture, fish guts and aquaria water).

**D, E, F:** Bacterial ASVs richness (**D**), Pielou’s evenness (**E**) and shannon (**F**) indices of the gut microbiota time-treatment groups. Letters refer to Benjamini-Hochberg (BH) adjusted Wilcoxon post-hoc test significance. Absence of letter refers to an absence of significant differences.

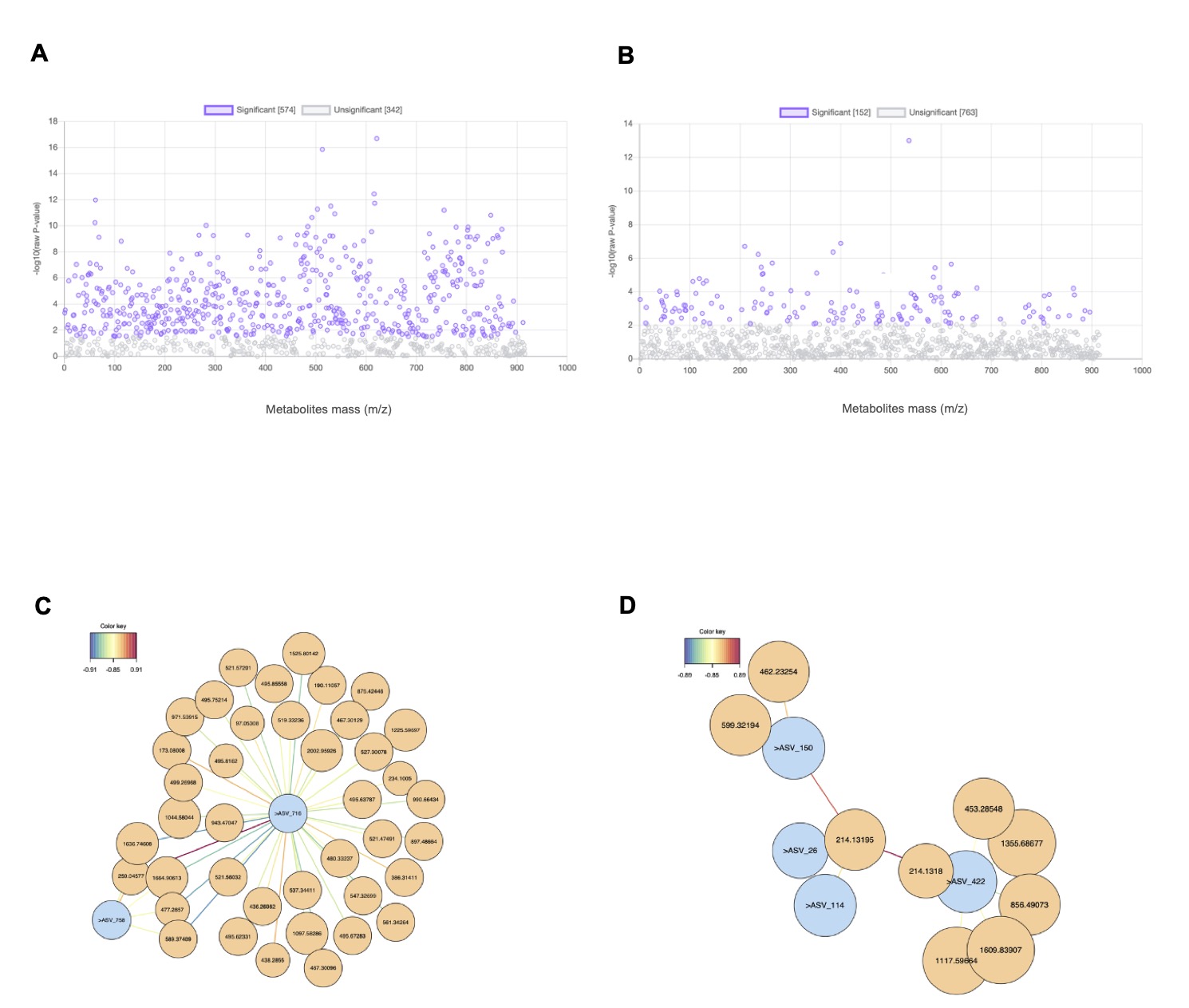

**Fig S2: Supplementary metabolome analysis figures.**

**A and B**: Gut metabolites differential abundances over time for the *M. aeruginosa* exposed (**A**) and control (**B**) treatments; Metabolites which are significantly differentially abundant between day 0 and 4 are colored in violet (color refer to ANOVA *p*<0.05).

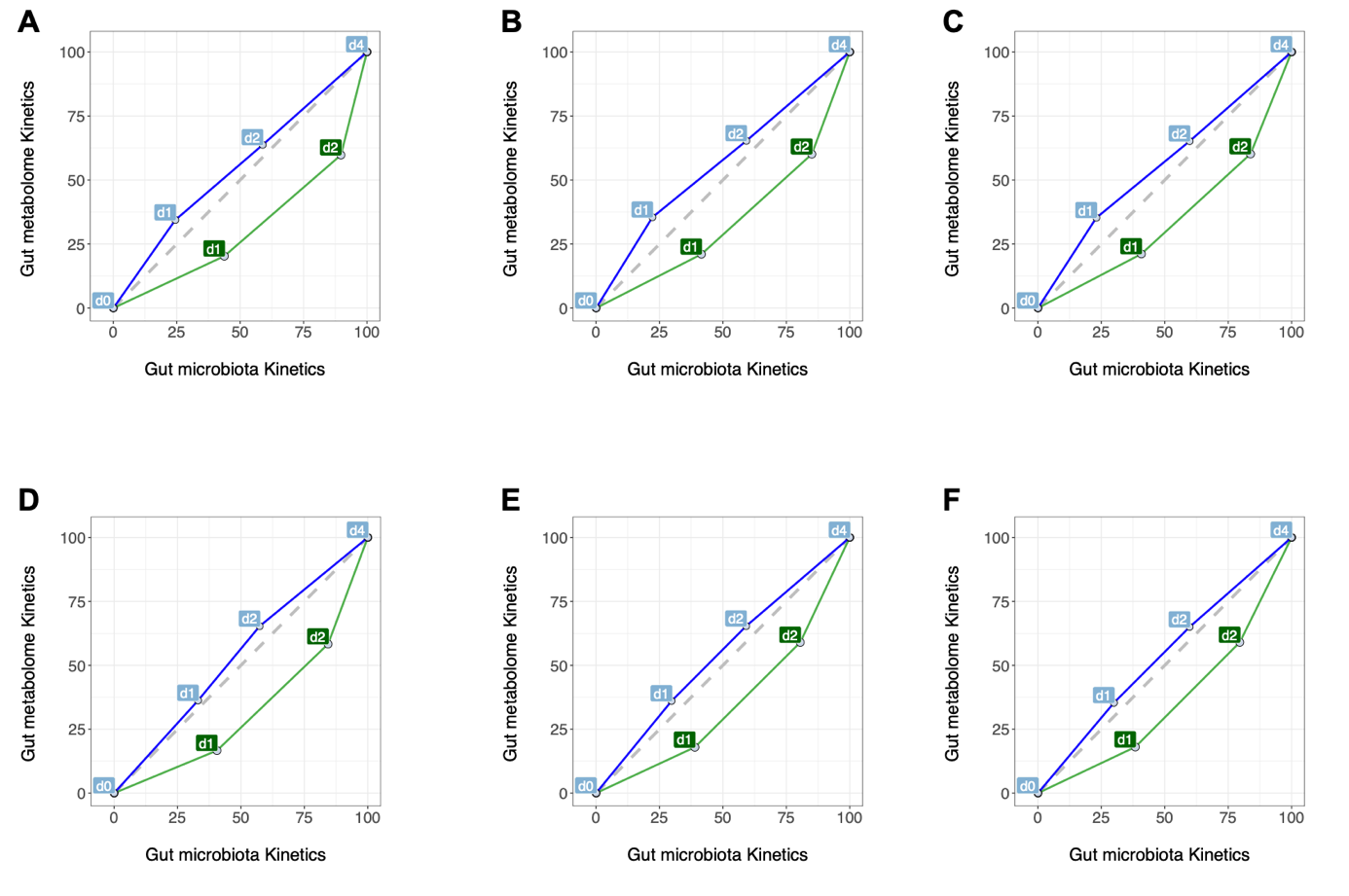

**Fig S3: Supplementary MOTA analysis figures.**

Trajectory of the composition change in microbiota versus metabolome expressed as percentages of the total trajectory achieved at day 0, day 1, day 2 and day 4 on each axis. Time_treatment group centroids coordinates were computed by mean (**A, B, C**) or median (**D, E, F**). The number of axes used were set to account for 70% (**A, D**), 90% (**B, E**) or 95% (**C, F**) of the total explained variance of the two datasets.
